# Closing the gap: Examining the impact of source habitat proximity on plant and soil microbial communities in post-mining spoil heap primary succession

**DOI:** 10.1101/2024.04.15.589563

**Authors:** Lenka Mészárošová, Eliška Kuťáková, Petr Kohout, Zuzana Münzbergová, Petr Baldrian

## Abstract

Revegetation of barren substrates is often determined by the composition and distance of the nearest plant community, serving as a source of colonising propagules. Whether such dispersal effect can be observed during the development of soil microbial communities, is not clear. In this study, we aimed to elucidate which factors structure plant and soil bacterial and fungal communities during primary succession on a limestone quarry spoil heap, focusing on the effect of distance to the adjoining xerophilous grassland.

We established a grid of 35 plots covering three successional stages – initial barren substrate, early successional community and late successional grassland ecosystem, the latter serving as the primary source of soil colonization. On these plots, we performed vegetation surveys of plant community composition and collected soil cores to analyse soil chemical properties and bacterial and fungal community composition.

The composition of early successional plant community was significantly affected by the proximity of the source late successional community, however, the effect weakened when the distance exceeded 20 m. Early successional microbial communities were structured mainly by the local plant community composition and soil chemical properties, with minimal contribution of the source community proximity.

These results show that on small spatial scales, species migration is an important determinant of plant community composition during primary succession while the establishment of soil microbial communities is not limited by dispersal and is primarily driven by local biotic and abiotic conditions.

## Introduction

Surface mining activities, such as limestone mining, degrade large areas of land, leaving behind a nutrient poor barren substrate. As such, they present unique ecological niches where primary succession during ecosystem recovery is steered jointly by microorganisms and plants. While primary succession has been mostly examined from a plant perspective, it is crucial to recognize that soil microbes act as the initial colonizers on newly exposed substrates (Nemergut et al., 2007; Harantová et al., 2017). They play a pivotal role in the accumulation and cycling of nutrients and soil organic matter before the onset of plant colonization (Kallenbach et al., 2016) and facilitate plant establishment (Schulz et al., 2013). This spontaneous succession is a cheap method to restore vegetation on degraded lands (Bradshaw, 1997), which often host endangered plant species adapted to sites with low nutrients and competition (Tropek et al., 2010). Therefore, spontaneous succession produces habitats with high conservation value (Prach et al., 2001, 2011; Novák and Konvička, 2006).

The process of plant community development is determined by factors such as propagule availability (Lichter, 2000; Lanta and Lepš, 2009), environmental conditions (Prach et al., 2007), and biotic interactions (Koffel et al., 2018). Propagule availability is closely linked to dispersal, and its importance for plant community assembly is especially high early in succession (Knappová and Münzbergová, 2015; Makoto and Wilson, 2019) when plants colonise empty niches and plant-plant interactions are minimal. Typically, most seeds disperse over a short distance from the parent plant with the number of seeds dispersing at longer distances dropping sharply (Cheplick, 2022). Accordingly, Novák and Konvička (2006) have shown that during the revegetation of abandoned quarries, the likelihood of successful establishment of a species depended on the size and proximity of its population in the surrounding undisturbed vegetation patches.

While microbes are generally less dispersal-limited than plants (Junker et al., 2021), dispersal still influences the assembly of soil microbial communities (J. Wang et al., 2022; Lemoine et al., 2023), especially fungi that seem to be substantially more dispersal limited than bacteria (Schmidt et al., 2014; Powell et al., 2015; Zhang et al., 2021). Soil microbes disperse through a variety of vectors, including animals, rain and wind (Huffman et al., 2013; Golan and Pringle, 2017), with patches of vegetation serving as sources of microbial propagules (Redondo et al., 2020), analogous to their role in plant colonisation. While information regarding the effect of distance source vegetation on microbial dispersal dynamics is sparse, especially for bacteria, it is conceivable that microbes exhibit dispersal patterns similar to plants (Barbour et al., 2023). Indeed, recent studies have shown that the abundance of fungal propagules decreases with increasing distance from potential spore sources (Peay et al., 2012; Grosdidier et al., 2018).

Plants further contribute to microbial community dispersal directly, introducing microbes into soil in and on seeds (Herrera Paredes and Lebeis, 2016), and indirectly, by attracting specific microbes via relatively long-distance (tens of cm) belowground chemical signalling (Bever et al., 2012; Schulz-Bohm et al., 2018). Additionally, plant growth modifies soil abiotic conditions and enriches soil with organic matter through litter inputs and rhizodeposition, largely in a plant-species specific manner (Urbanová et al., 2015; Steinauer et al., 2016; Kuťáková et al., 2020), potentially weakening environmental filters of microbial species establishment and selecting plant species-specific root or rhizosphere associated taxa (Mészárošová et al., 2024). Consequently, numerous studies have shown, that species composition of plant community is a strong structuring force of soil microbial community assembly (Leff et al., 2018; Hannula et al., 2019; Burrill et al., 2023).

This study aimed to investigate the effects of distance to the putative source habitat (between 10 and 50 meters approximately) on soil microbial community assembly and its associations with plants following the initial revegetation of a limestone quarry spoil deposit adjoining a late successional xerophilous grassland which served as a major source of both plant and microbial propagules. We hypothesised that i) the composition of the early successional (ES) plant community on the spoil heap will a visible distance gradient, i.e. that ES plots closer to the late successional (LS) grassland will have a plant community more similar to the LS community. ii) Consequently, we expected that plant community composition will impact on soil microbes, introducing a distance-related spatial pattern in microbial community composition, particularly in fungi that are known to be more responsive to vegetation composition than bacteria (Urbanová et al., 2015).

## Material and Methods

### Study site

The study was carried out on a limestone quarry spoil deposit and an adjacent species-rich dry calcareous grassland located in the Czech Karst Protected Landscape Area in the Central Bohemia, Czech Republic (49.96154 N, 14.16454 E – 49.96365 N, 14.16845 E). The surrounding hilly karstic landscape is characterized by relatively warm climate and mild winters (mean annual temperatures 8-9°C, mean annual precipitation 530 mm), the vegetation consists of thermophilic and xerothermic grasslands alongside deciduous forests and human settlements. Since 2009, the abandoned parts of the quarry have been gradually filled with clayey spoil from the deep layers of the quarry, and thus likely not containing any soil biota, and left to spontaneous revegetation. The oldest part of the spoil heap (ca. 150 x 100 m) represents an early primary successional habitat and is directly adjacent to a species-rich calcareous grassland with scattered trees, a late successional habitat, which serves as a source community for plant species colonizing the spoil heap (Kuťáková et al., 2020). Since 2005, the grassland has been grazed by sheep and goats.

### Soil and vegetation sampling

The sampling took place on June 25, 2015. To determine how the proximity of the source community influences the composition of plant and soil microbial communities during early succession, we established five sampling transects, each perpendicular to the grassland edge (Figure S1). At each transect, the first 1 m^2^ study plot was located within the late successional grassland, approx. 5 m from the grassland edge (LS) and another five points were located within the early successional (ES) spoil deposit area, 10 m (ES10), 20 m (ES20), 30 m (ES30), 40 m (ES40), and 50 m (ES50) from the first plot. Transects were parallel to each other and adjacent transects were separated by >10 meters (Figure S1). To get an idea how the microbial communities looked at the beginning of succession, we established additional six plots in the area with the most recent spoil deposition from 2014, representing the initial succession (IS) phase. These plots were located approximately 200 meters away from the last early successional plots. At each plot, we collected 5 soil cores (diameter of 35 mm), one in the centre and the other four in each corner of the plot. The samples were brought to the laboratory and processed within 24 h. Soil up to the depth of 10 cm from all cores from each plot were pooled, litter, stones and roots were removed, and the soil was sieved through a 5-mm sieve. The samples were freeze-dried, weighed and stored at −40°C for further analyses.

On each plot, vegetation composition was assessed by recording all species of vascular plants and visually estimating their percentage ground cover (Table S2). In order to capture all plant species present in the plots in their phenological optima, the vegetation sampling was conducted twice during the season of 2015, in the end of May and in the beginning of August, and the resulting dataset consisted of averaged values of the two observations. To assess which plant species are likely migrating from the adjacent grassland to the spoil deposit area, we used a detailed floristic survey performed in 2011 to a distance up to 100 m from the spoil heap (Kuťáková, unpublished data). The species found solely on the grassland but not in the other surrounding habitats were classified as grassland species. The nomenclature followed Kubát et al. (2002).

### Soil chemistry analyses

To characterize soil chemical properties, soil samples were analysed for pH (1:10 w/v soil-to-water ratio), available phosphorus (P_av_), total nitrogen (N_tot_) and total carbon (C_tot_). N_tot_, C_tot_ (using Carlo Erba Instrument NC 2500) and active pH (Zbíral, 2002) were measured by the Analytical laboratory of the Institute of Botany, Czech Academy of Sciences, Průhonice, Czech Republic. P_av_ was assessed in samples shaken with water (1:10, w:v) for 18 h, filtered through 0.22-μm membrane filter and measured with the malachite green method (Řezáčová et al., 2019). Soil moisture content was calculated based on the sample masses before and after freeze-drying. See Table S1 for the overview and soil properties of the study plots.

### DNA extraction & amplicon sequencing

Fungal and bacterial communities in soil and root samples were characterised by sequencing the internal transcribed spacer (ITS2) region and the V4 region of the 16S ribosomal RNA gene, respectively. DNA from each freeze-dried sample (350 mg of soil) was extracted in duplicates using the method of Miller modified by (Sagova-Mareckova et al., 2008). DNA extracts were purified using Geneclean Turbo Kit (Biogenic) following the manufactureŕs instructions, duplicates were pooled and stored in −20°C before further use. For the microbial community analysis, PCR amplification of the fungal ITS2 region from DNA was performed using barcoded primers fITS7 and ITS4 (Ihrmark et al., 2012). The V4 region of bacterial 16S rRNA was amplified using the barcoded primers 515F and 806R (Caporaso et al., 2012). PCR was performed in triplicate for each sample as described previously (Tláskal et al., 2016; Žifčáková et al., 2016). The resulting amplicons were purified, pooled, and libraries prepared with the TruSeq DNA PCR-Free Kit (Illumina) were sequenced in house on the Illumina MiSeq (2 × 250-base reads).

### Bioinformatic analysis

The amplicon sequencing data processing was done using the pipeline SEED 2.0.3 (Větrovský et al., 2018). Briefly, paired-end reads were merged using fastq-join (Aronesty, 2013) and fungal ITS2 region was extracted using ITSx 1.0.9 (Bengtsson-Palme et al., 2013). ITS sequences were then clustered at 97% similarity into Operational Taxonomic Units (OTUs) using UPARSE implemented in USEARCH7 (Edgar, 2013), detected chimeric sequences were deleted. The most abundant sequence was determined for each cluster, and its closest hit at a genus or species level was identified using blastn against the SILVA 138.1 (Quast et al., 2013) and UNITE 9.0 (Nilsson et al., 2019). Sequences identified as nonbacterial or nonfungal were discarded. Fungal guilds were determined based on the primary and secondary lifestyle in the FungalTraits database (Põlme et al., 2020). Sequence data have been deposited in the SRA under accession number PRJNA603889.

Illumina Miseq Sequencing yielded 596 904 bacterial and 433 179 fungal sequences. Bacterial sequences clustered into 15 122 OTUs, including 6 533 singletons; fungal sequences clustered into 13 191 OTUs, of which 8 268 were singletons. For microbial alpha diversity calculations, bacterial and fungal datasets including singletons were subsampled to the minimum number of reads (6506 for bacteria, 3390 for fungi). For further analyses, singletons and doubletons were filtered out and bacterial and fungal datasets were subsampled to 10 000 and 6 000 reads, respectively. We ended up with 336 757 bacterial sequences, clustering into 6 404 OTUs, and 202 720 fungal sequences clustering into 3 079 OTUs. Statistical analyses were performed on datasets containing only OTUs occurring with an abundance of 0.1% in at least three samples (3 072 bacterial OTUs, 96% of sequences; 1 775 fungal OTUs, 97% of sequences).

### Data analysis

Statistical analyses were performed using R software v. 4.1.0 (R Core Team and R Development Core Team, 2021), package *vegan* (Oksanen et al., 2022). The R-Code used in this article is available from the authors upon request.

Prior to statistical analyses, soil physicochemical variables except for pH (total soil N and C, available soil P, soil moisture) were log-transformed to normalize distribution and plant species cover data and bacterial and fungal OTU abundances were Hellinger-transformed. One of the fungal LS samples was dropped due to due to poor sequencing performance.

We used linear regression to investigate how individual soil chemical characteristics, plant species richness and cover, and soil microbial species richness change with distance from the source grassland community. To assess variation in beta diversities among microbial groups and plots, we measured the multivariate homogeneity of group dispersions using the function betadisper based on Bray-Curtis dissimilarities (Anderson et al., 2006); the results were visualised as mean distance-to-centroid values. As we wanted to detect the effect of source community proximity on microbial community assembly on the spatial scale, we compared the community composition dissimilarity (Bray-Curtis) of sample pairs within each transect considering their distances. Compositional dissimilarities within the ES distance classes and between ES and IS transect plots were calculated analogously. The overall correlation between bacterial and fungal communities was calculated on full Bray-Curtis dissimilarity matrices using Mantel test.

To test if soil chemical properties, plant community composition and soil microbial community composition differed across transects, we performed permutational multivariate analysis of variance (PERMANOVA) on respective datasets. Compositional differences of communities between pairs of ES distance classes were tested using pairwise PERMANOVA with Benjamini-Hochberg correction for multiple testing. We used PCA and NMDS to visualise the compositional differences among samples, using the *envfit* function to fit the environmental variables onto the ordinations. In the case of plants, we depicted only those plant species that contributed more than 30% of variation (P < 0.05), as determined by *envfit*.

To assess the relative importance of space, soil chemistry, and plant community composition for structuring soil microbial communities during early succession, we performed the variation partitioning analysis of ES microbial communities (Legendre et al., 2005) using the function *varpart*. Plant community data were represented by the scores of the NMDS axes. The spatial predictors were derived from the GPS coordinates of individual samples using the principal coordinates of neighbourhood matrix (PCNM) method (Borcard and Legendre, 2002). To select the most relevant predictors of microbial data variation, independent double-stop forward selections (*adespatial*) (Dray et al., 2023) of soil chemical variables and PCNM vectors were carried out prior to the variation partitioning. To disentangle the effects of distance and distance-unrelated spatial variability, we detrended bacterial, fungal and plant community data to remove the distance aspect represented as linear and second order polynomial terms prior to forward selection of relevant PCNM vectors.

## Results and Discussion

### Soil chemistry & Plant community composition

Soil chemistry differed by habitat, with soil nutrient content increasing and soil pH decreasing with habitat successional stage. The early successional plots (ES) were more similar to the initial (IS) than to the late successional (LS) soil and did not show significant differences among distance classes (Fig. 1). However, soil pH of the ES plots significantly increased with distance from the late successional habitat (R^2^adj = 0.18, P < 0.05, Fig. 1).

**Figure 1.**
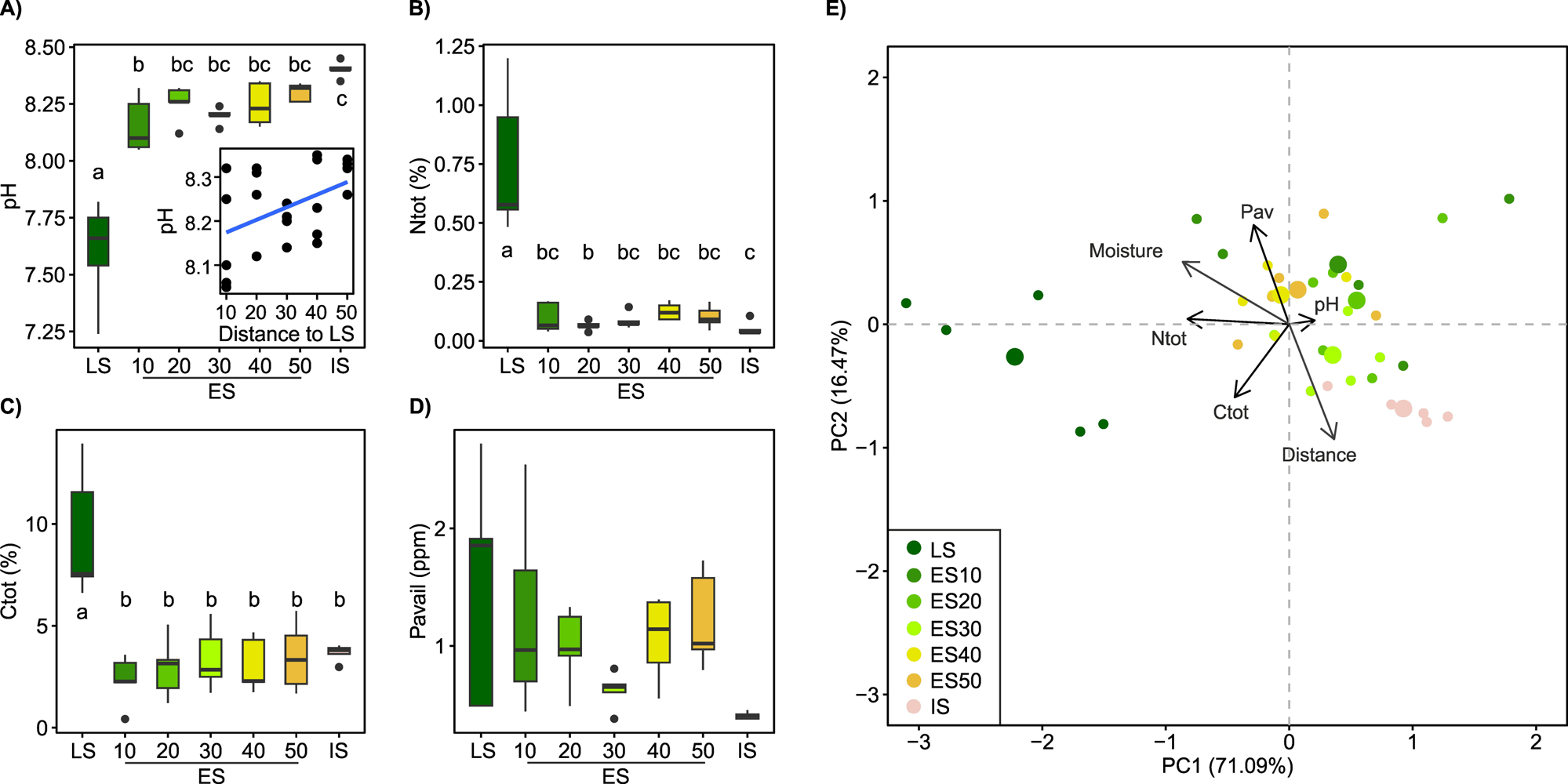
Soil properties, pH, total N, total C, and available P, across a successional grassland sequence. In (**A-D**), treatments with different letters are significantly different (P< 0.05, ANOVA, Tukeýs test). The scatterplot within the pH boxplot depicts the correlation between pH and distance to the LS for the ES plots. (**E**) The principal component analysis (PCA) biplot shows clustering of samples based on their soil chemical properties, the “Distance” vector represents the distance to the LS and was added to the ordination as a supplementary variable using envfit. LS – late successional plots, ES – early successional plots, numbers denote distance (in m) to the LS, IS – initial soil.

Plant species richness and vegetation cover of the ES plots were not significantly different from those of the LS community, but both tended to decrease with distance from the LS (Fig. 2a, b). Plant species richness across the ES transects was significantly negatively correlated with soil N_tot_ and P_av_ (Fig. S2). The grassland plant species cover significantly decreased with distance to the LS (R^2^adj = 0.43, P < 0.001, Fig. 2c, d). Plant community composition in ES significantly differed not only from LS plant community but also among ES distance classes, with ES10 being significantly different from ES30-50 and ES20 falling in between (Fig. 2e). Linear regression analysis further showed that the similarity of ES and LS plant communities significantly decreased with distance (Fig. 3c) while plant cover showed marginally significant decrease (Fig. S2). A significant decrease of similarity with distance was also observed among ES communities (Fig. 3f). ES plant community variation exhibited strong spatial structure. The most significant factor influencing plant community composition across the ES plots was the distance to the LS, accounting for over 11% of the community variation. Additionally, spatial heterogeneity unrelated to distance explained another 5% of the community variation (Fig. S3). These results are consistent with our first hypothesis, confirming the decisive role of source community proximity in the assembly of ES plant communities. In line with the findings of Novák and Konvička (2006), who demonstrated distance-dependent differences in successional trajectories and establishment success of source grassland species in abandoned basalt quarries, we observed a tipping point in ES plant community assembly at a distance of 30 meters from LS (Fig. 2e). The plant community composition of the more proximal ES plots was more similar to the LS and tended to contain more LS grassland species than the ES plots located further away.

**Figure 2.**
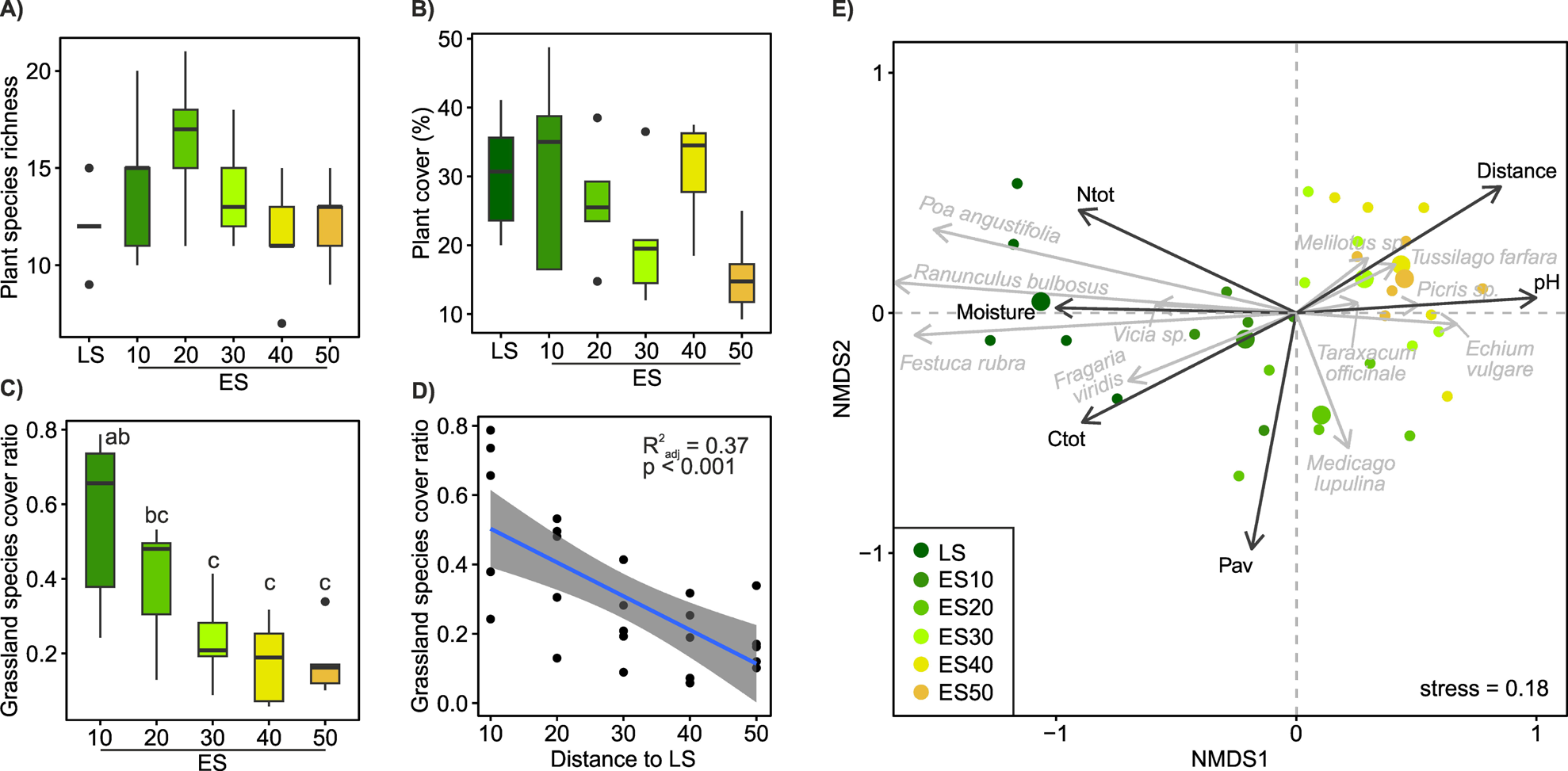
Plant community composition across a successional grassland sequence. (**A-D**) Plant community properties. Treatments with different letters are significantly different (P< 0.05, ANOVA, Tukeýs test). The scatter plot illustrates the relationship between the share of grassland plant species cover in ES plots and their distance from the LS plots. (**E**) Beta-diversity of the plant community. Non-metric multidimensional scaling (NMDS) ordination based on Bray-Curtis distances calculated on Hellinger-transformed plant species cover data (grey arrows), environmental variables significantly correlated (P < 0.05) with ordination axes are shown as vectors (black arrows).

**Figure 3.**
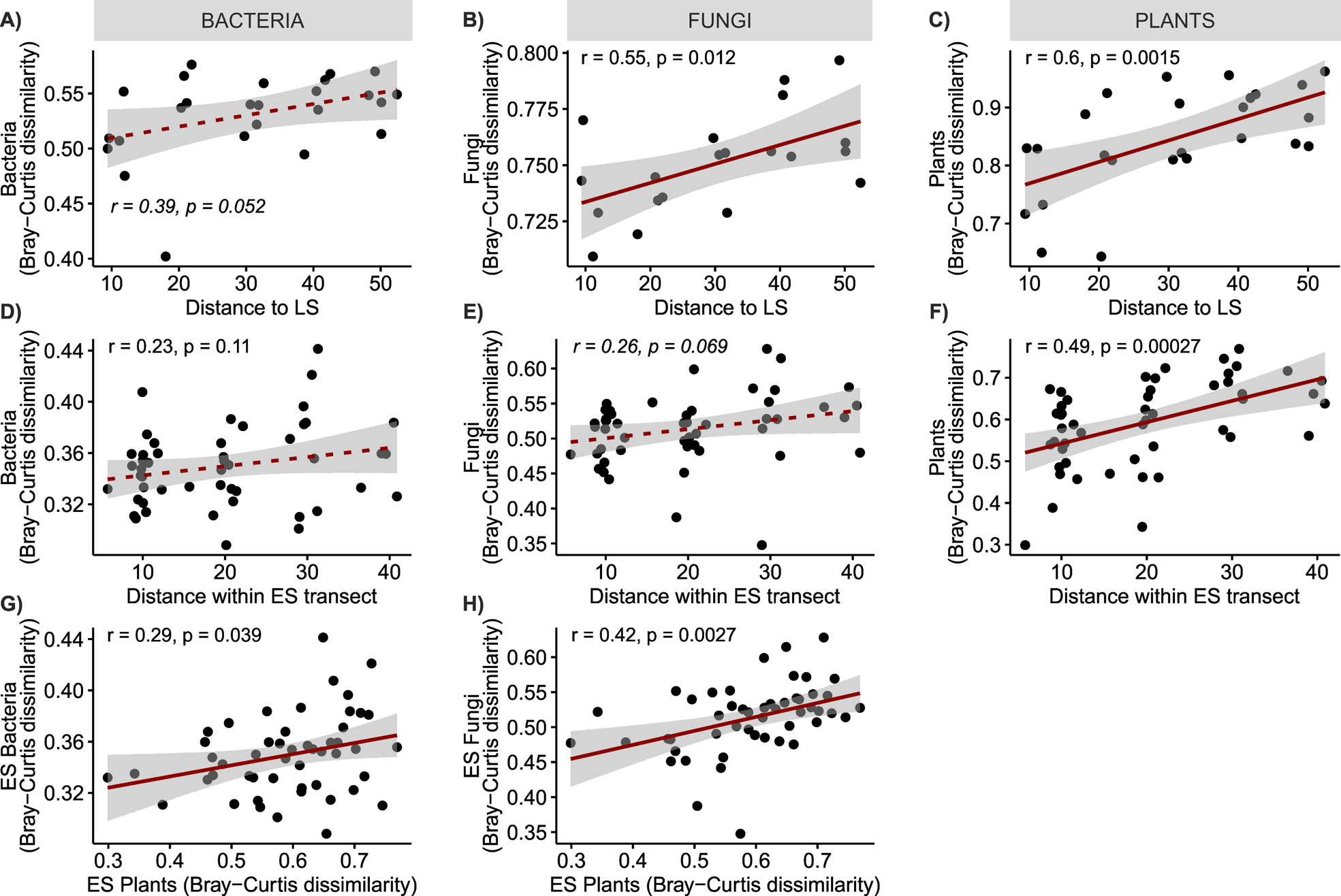
Linear regressions of (**A-C**) the dissimilarity of ES plot communities from the LS community and distance to LS, (**D-F**) dissimilarity of ES plant communities and distance between ES plots, and (**G-H**) similarity of microbial and plant community composition. Solid lines denote significant relationships at P < 0.05, dashed lines denote non-significant relationship.

### Microbial community richness and composition

The differences in taxonomic composition of both bacterial and fungal communities were especially pronounced at finer taxonomic levels (Fig. S4). Bacterial communities exhibited higher overall evenness compared to fungal communities, and both microbial groups exhibited significant differences in community evenness across the three successional habitats, with the highest evenness observed in the ES communities. Fungal IS community was particularly uneven with almost 70% of sequences belonging to top five OTUs, whereas the LS and ES communities were more balanced (Fig. S4c, d).

Both bacterial and fungal OTU richness tended to be higher in the ES soil than in the LS or IS soil (Fig. 4a, b, d, e). Bacterial richness in ES was influenced neither by distance to the LS transect, nor by any of the soil properties or plant species richness and/or cover. Contrastingly, ES fungal richness was significantly positively correlated with plant species richness and soil moisture (Fig. S2). The relationship between plant and soil microbial species richness has been extensively studied, with varied results. Some studies in grasslands have reported a positive correlation between plant and soil fungal (Chen et al., 2017; Dassen et al., 2017; Yang et al., 2017) or bacterial (Yang et al., 2018) species richness, while others found no significant relationship (Prober et al., 2015; Navrátilová et al., 2018; Guasconi et al., 2023). Our results suggest a stronger link between plant and soil fungal richness compared to bacterial richness, consistent with the findings by Wang et al. (2022).

**Figure 4.**
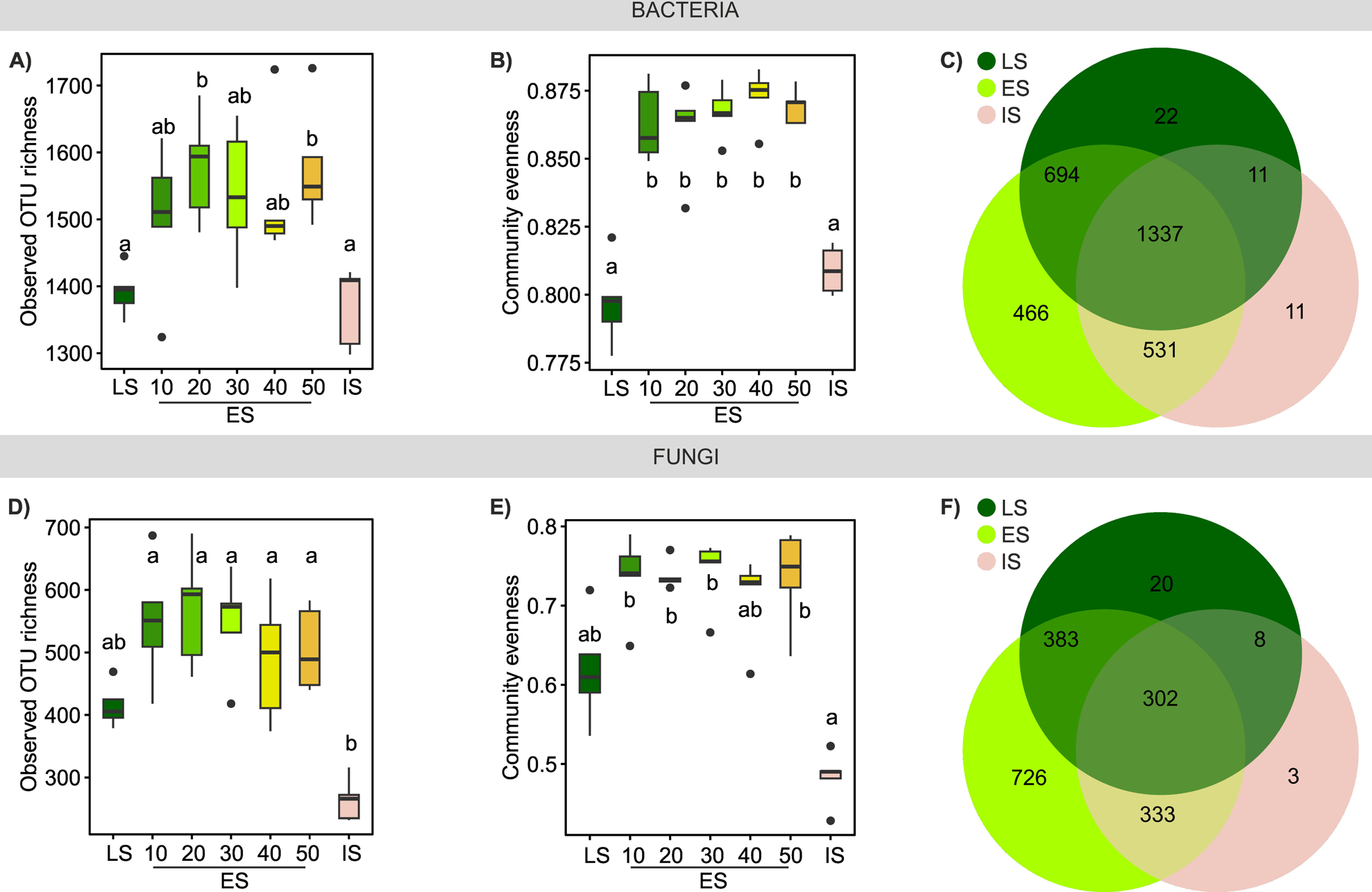
Microbial richness and evenness across a successional grassland sequence. (**A, D**) Observed OTU richness and (**B, E**) community evenness of bacterial and fungal communities; treatments with different letters are significantly different (P< 0.05, ANOVA, Tukeýs test). (**C, F**) Venn diagrams illustrate the number of OTUs specific to each habitat or shared among them.

Plant species richness could influence fungal richness in several ways. Firstly, higher plant species diversity affects the quantity and chemical diversity of plant litter (Schnitzer et al., 2011; Santonja et al., 2017), modulates metabolite production of individual plant species (Scherling et al., 2010), and enhances the amount and diversity of plant exudates (Steinauer et al., 2016; Eisenhauer et al., 2017). With fungi being the main decomposers of recalcitrant plant-derived organic matter (López-Mondéjar et al., 2018) and receivers of a substantial portion of recent plant photosynthate carbon (Hawkins et al., 2023), the enhanced diversity of plant-derived nutrients may directly influence the diversity of soil fungal communities (Broeckling et al., 2008; Cline and Zak, 2015). Secondly, higher plant species richness has been associated with greater diversity in communities of fungal plant pathogens (Liu et al., 2021) and AMF (Hiiesalu et al., 2014), fungal guilds particularly abundant in the ES community (Fig. S4e). The positive correlation between fungal species richness and soil moisture suggests a potential role for microclimate, possibly through increased shading of more species-rich ES plots (Fischer et al., 2019). Alternatively, the earlier attainment of higher plant cover in ES plots closer to the grassland may also contribute to this correlation. This early plant cover could lead to the accumulation of more organic matter, subsequently increasing water retention potential.

Bacterial and fungal community similarities across the three successional habitats were substantially correlated (Fig. S5a), indicating that their assembly is governed by similar factors; however, the beta diversity of bacterial communities was significantly lower than that of fungal communities (Fig. S5b). While both bacterial and fungal communities significantly differed by habitat (PERMANOVA R^2^adj = 0.46, P < 0.001 and R^2^adj = 0.31, P < 0.001, respectively, Fig. 5a, b), bacterial taxa were more ubiquitous than those of fungi, resulting in a larger community overlap across the habitats, i.e. higher proportion of bacterial OTUs shared among the ES, LS, and IS habitats compared to fungal OTUs. The proportion of OTUs unique to the IS habitat was notably low for both bacteria and fungi (Fig. 4c, f).

**Figure 5.**
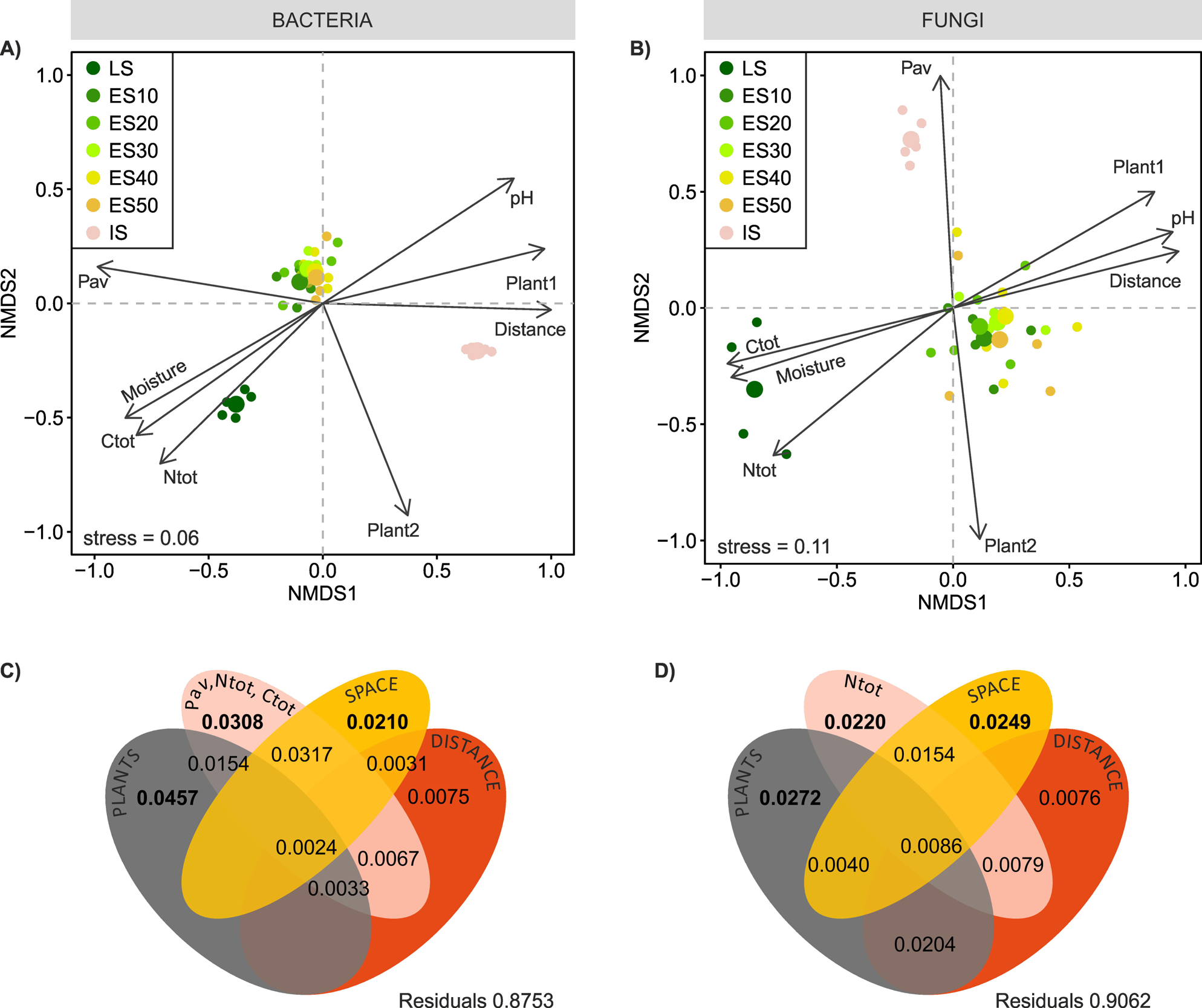
Bacterial and fungal community composition across a successional grassland sequence. (**A, B**) Beta-diversity of the microbial communities. Non-metric multidimensional scaling (NMDS) ordination based on Bray-Curtis distances calculated on Hellinger-transformed OTU abundance data, environmental variables were fitted onto the ordinations as vectors using envfit. Plant1, 2 – scores of NMDS axes 1 and 2 (**C, D**) Partitioning of variance in ES microbial community composition among plant community composition, soil chemistry, spatial variables, and their joint effects as predictors. For each testable fraction, an adjusted R2 is given, and significance (P < 0.05) is indicated in bold.

We expected to observe a clear distance gradient in microbial ES community composition, however, our assumptions were only partially met. Microbial communities across ES transects were rather homogeneous, with bacteria more so than fungi, and their composition did not exhibit any apparent distance-related patterns akin to those observed in plants. These results concur with those of García de León et al. (2016) who found no clear effect of presumed propagule source distance on the small-scale dispersal (tens of meters) of AMF during early succession, Bacterial ES community composition was more similar to the LS than IS community (average pairwise community dissimilarity was 0.53 ± 0.007 and 0.60 ± 0.008, respectively, P < 0.001), while fungal ES community resembled more the IS than the LS community (average pairwise community dissimilarity was 0.70 ± 0.015 and 0.75 ± 0.005, respectively, P < 0.01). Both bacterial and fungal ES communities became progressively more dissimilar to the LS community with increasing distance (Fig. 3a, b). Additionally, both bacterial and fungal ES community dissimilarity was significantly positively correlated with plant ES community dissimilarity (Fig. 3g, h). However, in both cases, the relationships were more pronounced for fungi than bacteria. The prominent role of plant community composition in structuring the ES soil microbial communities was confirmed by the variation partitioning. The analysis explained 9.5 and 12.6% of the variation in ES fungal and bacterial community composition, respectively, and identified the ES plant community composition as the most important determinant of variation in both the bacterial and the fungal communities, which is in line with previous studies on microbial succession (Knelman et al., 2012; Harantová et al., 2017). The second strongest driver of microbial community variation was soil nutrient content, followed by distance-unrelated spatial structuring. Bacteria in particular appeared sensitive to spatial gradients of soil nutrients evidenced by the shared effect of space and soil chemistry. The distance to the LS *per se* did not directly affect bacterial or fungal community composition. Instead, its impact on fungal, but not bacterial, community composition manifested indirectly through shared effects with plants, reflecting spatial gradients within the ES plant community composition (Fig. 5c, d).

In any case, the spatial effects were rather weak, and our results suggest that during primary succession on small spatial scales (tens of meters), the development of soil microbial communities is likely constrained more by local abiotic conditions than by dispersal limitations. Alternatively, the non-existence of a readily visible distance gradient in the composition of ES microbial communities might be due to the fact that the succession on the spoil heap was in its very early stage, with only 6 years of development. Our previous study on succession (Harantová et al., 2017) showed that fungal communities started to diverge later, with more profound changes in vegetation composition, and changes in bacterial community composition were even less evident. In the present study, taxonomic profiles of ES bacterial and fungal communities fall somewhere between LS and IS communities, but they more closely resemble the IS soil. However, given that ES plant community composition was the primary determinant of the structure of ES microbial communities and the covariation of ES plant and fungal community dissimilarities, it can be envisaged that soil microbes, in particular fungi, will follow changes in plant community composition and with time, the effect of source community proximity might become more apparent. Conversely, the grassland developing at ES may become progressively more homogenous across ES distance classes as plants overcome dispersal limitations.

It should be noted that the plant effect is stronger in root microbiomes than in surrounding soil (Mészárošová et al., 2024). One can thus not rule out that the differences in plant communities in the ES, while not affecting soil microbes, might possibly be visible in the root microbiome.

## Conclusion

Source habitat proximity was the main structuring force of early successional plant community composition, while having only minimal influence over the early successional microbial communities. Soil microbial community variation was determined primarily by early successional plant community composition and soil nutrient content, indicating local conditions play a more significant role than dispersal limitations. Even though bacterial and fungal early successional community assembly was governed by similar factors, there were observable differences – fungal community composition appeared more sensitive to plant community variation, whereas bacterial community was more affected more by edaphic properties.

## Acknowledgements

The study was supported by the Czech Science Foundation (project 15-11635S).

This work was supported by the Czech Science Foundation (21-17749S) and by the Ministry of Education, Youth and Sports of the Czech Republic (CZ.02.01.01/00/22_–08/000463– - AdAgriF - Advanced methods of greenhouse gases emission reduction and sequestration in agriculture and forest landscape for climate change mitigation).

## Supplementary Tables

Table S1. Details on study plots

Table S2 Floristic composition of study plots

## Notes

https://www.ebi.ac.uk/ena/browser/view/PRJNA603889

### Competing Interest Statement

The authors have declared no competing interest.

