## Supplementary Figures 1-5 for "Closing the gap: Examining the impact of source habitat proximity on plant and soil microbial communities in post-mining spoil heap primary succession"

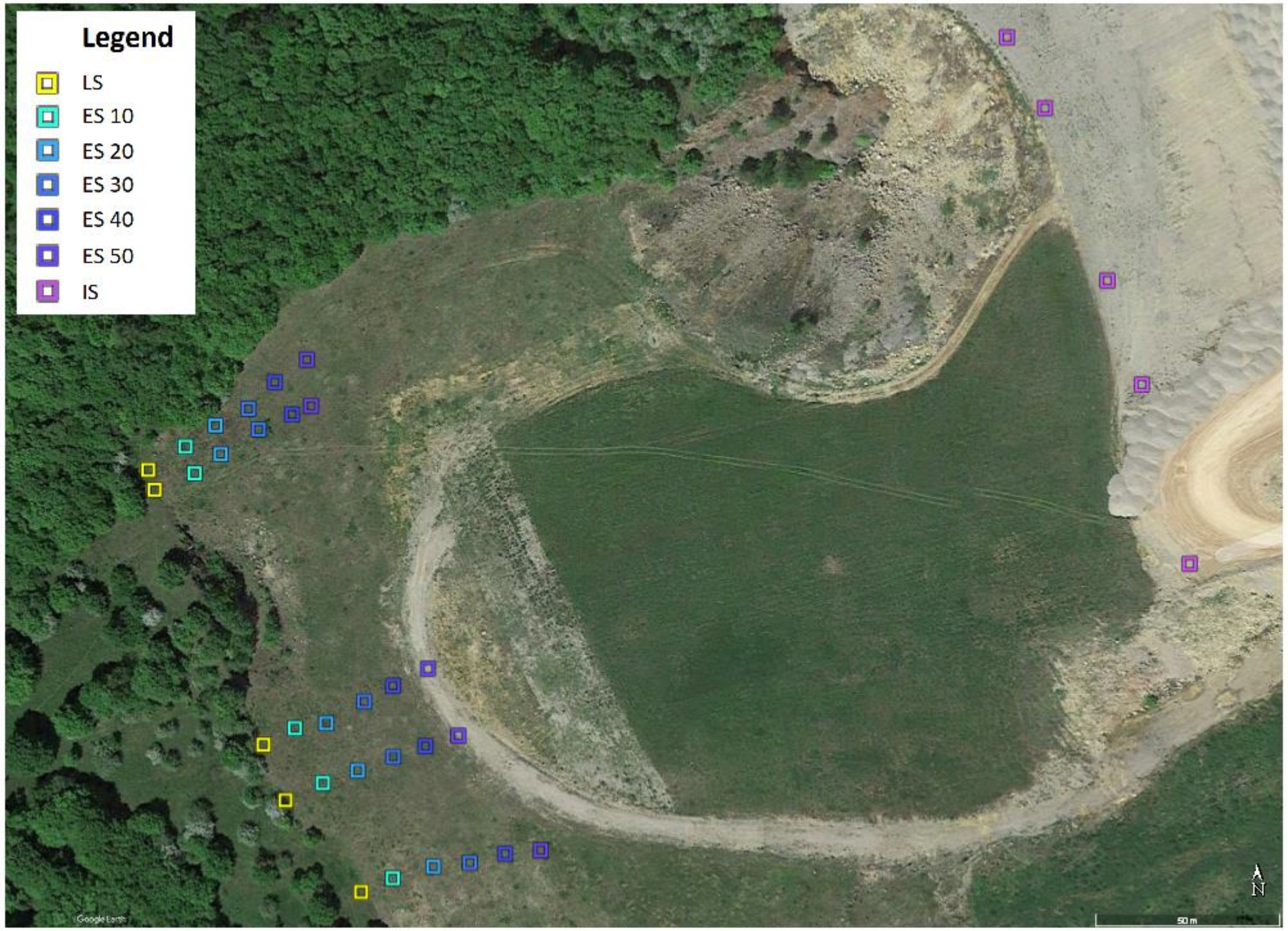


**Figure S1.** Location of the sampling plots within the study area of the successional grassland sequence.

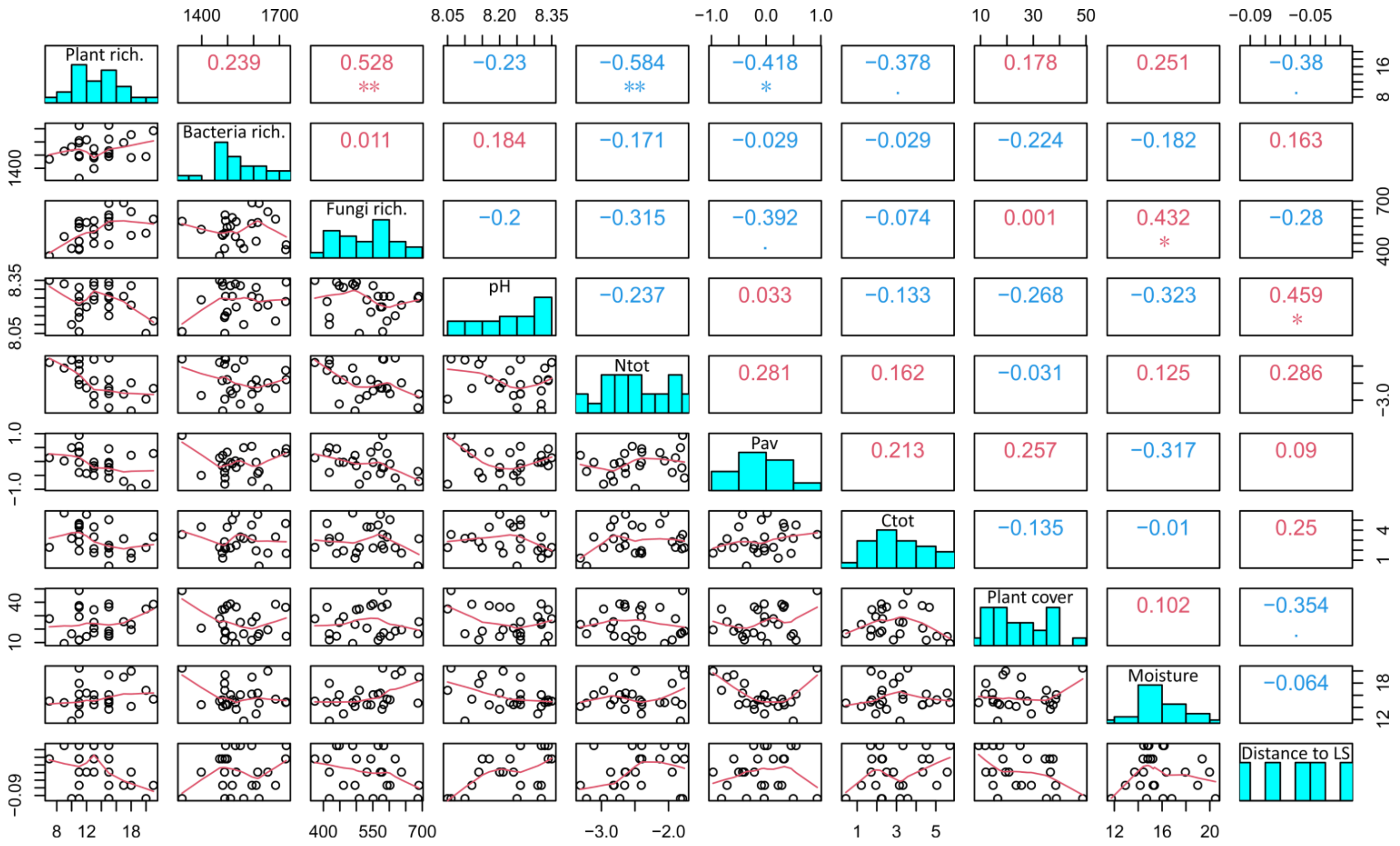


**Figure S2.** Correlations of environmental variables in a successional grassland sequence. Pairwise scatter plot matrix (lower panel), frequency distribution histogram (main diagonal), and Spearman correlation coefficients (upper panel) of plant and microbial species richness and environmental variables in the ES plots.
* P < 0.05; ** P < 0.01; *** P < 0.001

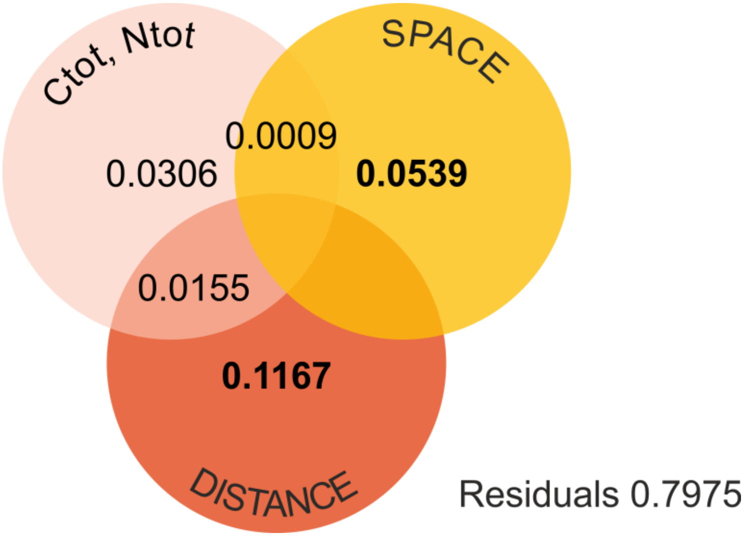


**Figure S3.** Partitioning of variance in ES plant community composition among soil chemistry, distance-unrelated spatial variables, and distance to LS, and their joint effects as predictors. For each testable fraction, an adjusted R2 is given, and significance ($P < 0.05$) is indicated in bold.

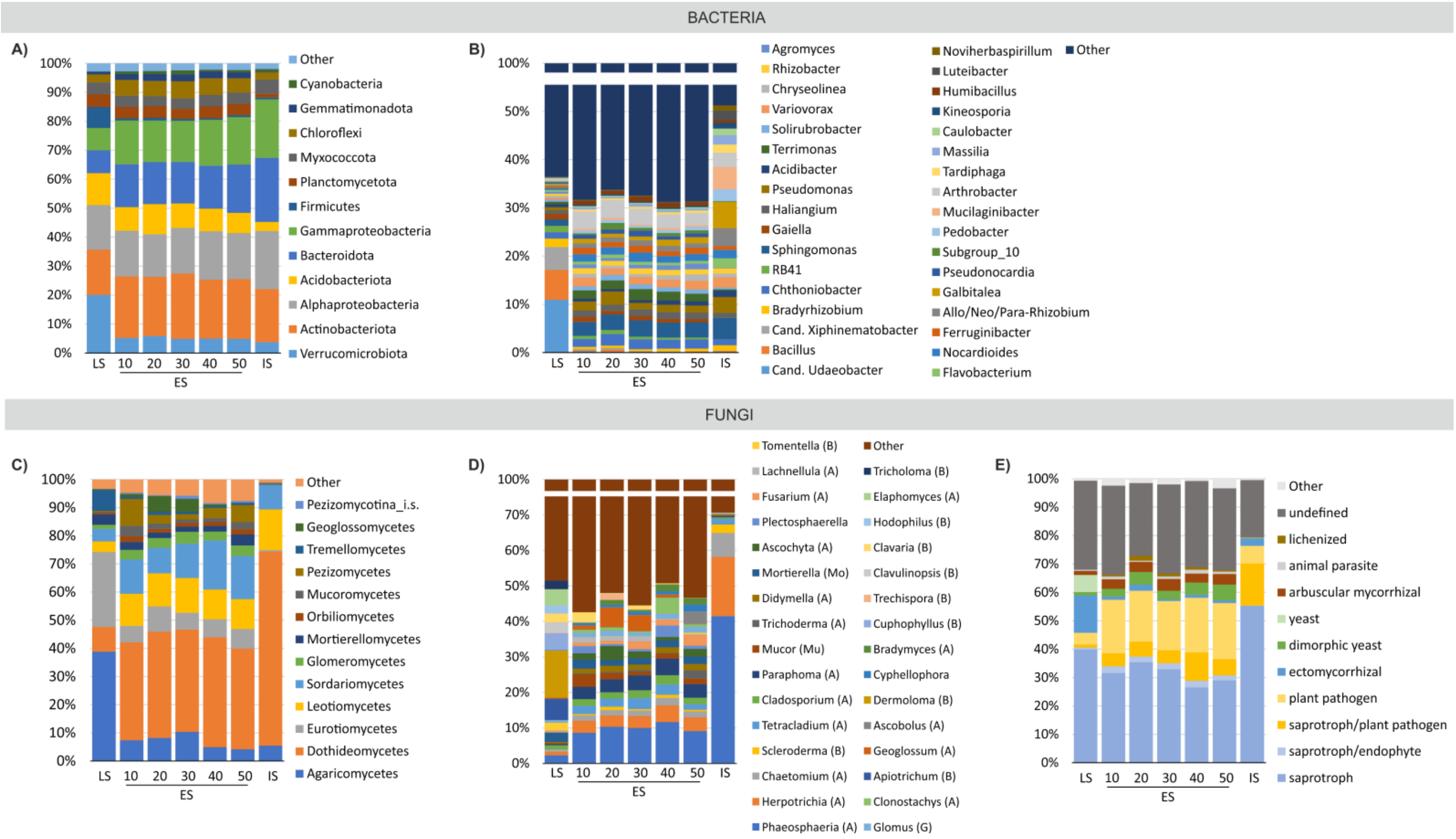


**Figure S4.** Taxonomic profiles of microbial communities in a successional grassland sequence. Bar graphs illustrate the taxonomic composition of (**A, B**) bacterial and (**C, D**) fungal communities and (**E**) fungal ecological guilds across plot classes (n=5).

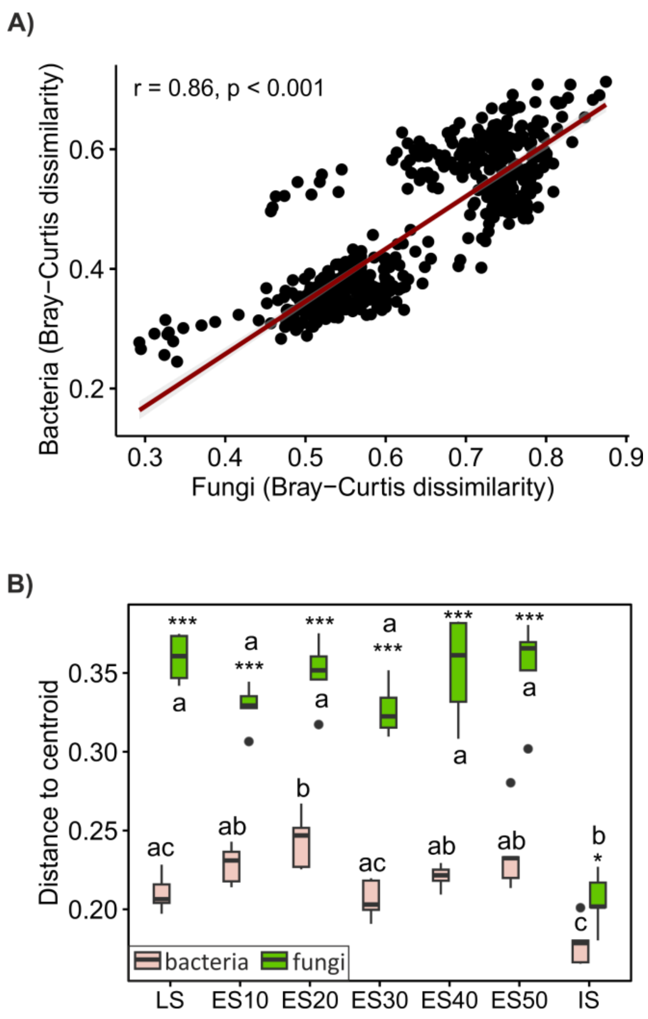


**Figure S5. A**) Linear regression analysis depicting the relationship between pairwise similarities of bacterial and fungal communities based on Bray-Curtis dissimilarities. **B**) Analysis of within treatment variation in microbial community composition. Betadisper analysis comparing bacterial and fungal multivariate dispersion across treatments. Different letters indicate significant differences ($P < 0.05$) among plot classes within each microbial group, asterisks denote significant differences between microbial groups. * $P<0.05$; *** $P < 0.001$.
